# De novo design of ligand binding and sensing with a physics based generative approach

**DOI:** 10.64898/2026.07.13.738243

**Authors:** Yanzhe Zhang, Yitao Ke, Rui Zhi, Qihan Jin, Yiqing Feng, Chentong Wang, Minchao Fang, Jinyang Liao, Dachuan Chen, Jiao Liu, Longxing Cao

**Affiliations:** State Key Laboratory of Gene Expression, School of Life Sciences, Westlake University, Hangzhou 310024, China; Artificial Intelligence Drug Design Core Laboratory, Westlake Laboratory of Life Sciences and Biomedicine, Hangzhou, Zhejiang, 310024, China

## Abstract

The de novo design of ligand-binding proteins has tremendous potential to revolutionize biosensor technology, yet converting these designs into functional sensors remains a major challenge due to the need for ligand-induced conformational changes or modulation of protein–protein interactions. Here, we introduce a physics-based generative approach for the de novo creation of proteins that bind small molecules and metal ions. Our method achieves customizable ligand-binding pocket formation in parallel with simulated protein folding, allowing for precise architectural control of the protein–ligand complex and facilitating the development of biosensors based on either ligand-triggered protein reassociation via split-protein reassembly or ligand-induced protein folding. We demonstrate the versatility of our computational method through successful designs targeting five small molecules, including the very small neurotransmitters serotonin and dopamine, and two metal ions. Biophysical characterization confirmed correct ligand binding, and crystal structures closely matched computational models. We demonstrated the biosensor engineering potential of these designs by constructing serotonin and dopamine sensors using a split protein strategy and explored several approaches to enhance sensor activity. Additionally, we developed a zinc sensor through a zinc-induced protein folding mechanism. Overall, our physics-based generative approach provides a robust framework for the de novo design of ligand-binding proteins, opening new avenues for the development of ligand-responsive biosensors.

## Introduction

Biosensors are indispensable analytical tools that transduce a biological recognition event into a quantifiable signal, enabling the sensitive and selective detection of diverse analytes, including small molecules and metal ions^1,2,3,4^. Most biosensors consist of a sensing module that switches configuration or modulates the association of protein domains upon specific target binding, and a transducer—often a fluorescent reporter—that converts this event into a measurable signal^5,6^. Despite their essential roles in bioprocess monitoring, clinical diagnostics, and environmental sensing, the development of new biosensors remains a considerable challenge^7,8^. A widely adopted strategy involves integrating ligand-responsive domains with cpGFP-based systems for analyte detection^5,7,9^. However, these approaches largely depend on naturally occurring ligand sensing domains, whose conformational dynamics and specificity are often difficult to re-engineer^4^. Consequently, expanding the repertoire of biosensors to detect novel analytes typically demands extensive rounds of screening and optimization, which may slow progress in the field^10^.

Rapid advances in protein design have enabled promising strategies for constructing small molecule- and metal ion-binding proteins, demonstrating considerable potential for the development of new biosensors^11,12,13,14,15,16^. Traditionally, physics-based approaches to de novo ligand binder design involve generating large libraries of pocket containing protein scaffolds, followed by computational docking and sequence design to identify favorable ligand interactions^11,17,18,19,11,16^. However, the vast chemical diversity of small molecules means that scaffold libraries rarely contain structures perfectly suited for every ligand. This challenge is analogous to in silico virtual screening in drug discovery, where extremely large chemical libraries are commonly required to identify a suitable candidate^20,21^. Recently, deep learning-based generative models have enabled the design of proteins with promising ligand-binding capabilities^22,23,24,25,26^. Nevertheless, most deep learning based approaches lack interpretability and offer limited control over specific protein–ligand interaction features. Furthermore, these designs typically emphasize static, high-affinity binding, and the resulting proteins seldom undergo significant ligand-induced conformational changes, thus limiting their direct applicability as biosensor modules. Only a few studies have demonstrated that de novo designed binders can be converted into functional biosensors^11,12,13,14,15^, typically by splitting the protein to facilitate ligand-induced reassembly^14,16^,, or by creating new binders that specifically recognize ligand-exposed surfaces on protein–ligand complexes^15,27^. Consequently, there is a critical need for protein design methodologies that generate small molecule and metal ion binders with structural attributes directly amenable to biosensor engineering. Advancements in this area would greatly accelerate the development of robust, specific, and modular biosensors for the precise monitoring of diverse biological processes.

### Design pipeline

We aimed to develop a robust and generalizable de novo protein design pipeline capable of accommodating the structural and chemical diversity of both small molecules and metal ions (**Fig. 1**). Our objective was to generate protein scaffolds that are readily adaptable for biosensor development, either by facilitating efficient splitting into two fragments—enabling ligand binding to drive fragment association—or by enabling ligand binding to be tightly coupled to protein folding, thereby inducing substantial conformational changes. Realizing these functionalities requires precise control over the overall topology of the protein scaffold to support protein splitting, and/or a careful balance between protein folding stability and protein–ligand interactions to allow for effective ligand-induced conformational changes. To address the diverse structural and chemical requirements of different ligands, we developed a bottom-up, physics-based generative design pipeline that simultaneously constructs protein backbones and optimizes protein–ligand interactions.

**Fig. 1:**
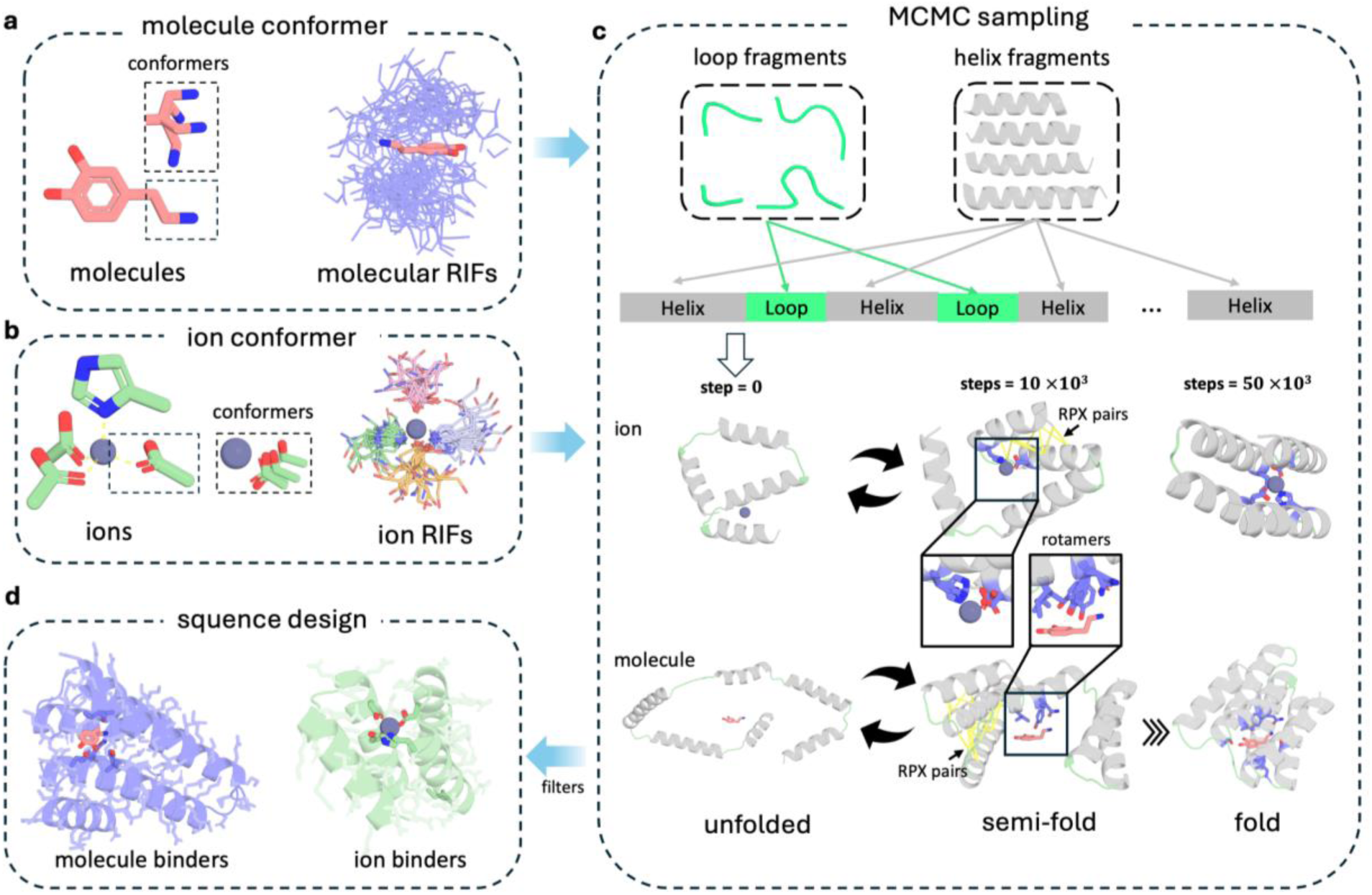
Pipeline of the physical-based generative method for designing ligand binders. **a**, Generation of small molecule Rotamer Interaction Field (RIF) for explicit modeling of protein–small molecule interactions. Small molecule conformers were sampled, clustered, and refined through quantum optimization, and RIFs were generated from these optimized conformers using RIFGen. **b**, Generation of metal ion coordination RIF. For each metal ion, the coordination geometry and residue types were defined, and the coordination rotamer at each coordination site was systematically sampled. **c**, Protein generation using a physics-based generative approach. A fragment assembly method was employed to build proteins. RPX and RIF scores guided protein folding and ligand binding during Monte Carlo simulated annealing, with the folding process informed by protein–ligand interactions (RIF score) and internal protein stability (RPX score), progressing from unfolded to semi-folded and folded states. The folding root, which remains fixed during fragment assembly, is initially selected at random. During sampling, it can be switched to other RIF-anchoring residues to comprehensively explore the relative positions of the protein and the ligand (**Supplementary Fig. 1**). **d**, Sequence design and final filtering. Well-packed scaffolds were selected for sequence optimization, and top-scoring designs were chosen for experimental characterization.

For small molecules, our design strategy begins with the generation of representative conformer ensembles for each ligand^28,29^. We then apply the rotamer interaction field (RIF) methodology^19,30^ to systematically enumerate energetically favorable side-chain interactions with the target ligand, yielding estimates of interaction energies and a map of critical contact residues (**Fig. 1a**). Backbone positions of the RIF residues serve as anchor points, guiding backbone construction to create protein scaffolds that are tightly wrapped around the ligand, thereby maximizing both shape and chemical complementarity. Privileged interactions, such as key hydrogen bonds, could be enforced or upweighted during structure generation to enhance binding affinity and specificity.

Metal ion binding requires precise combinations of coordinating residues arranged in well-defined geometries^31^. For example, zinc ions typically favor tetrahedral coordination involving histidine and aspartate/glutamate residues, while nickel ions generally adopt an octahedral coordination geometry. we constructed a coordination residue RIF table by systematically sampling all permissible coordinating residues based on the preferred residue type and the metal ion’s coordination geometry (**Fig. 1b**). Each entry in the coordination RIF is labeled with its intended coordination site, enabling fine control over the formation of complete or partial metal coordination spheres during protein backbone construction.

Rather than relying on the docking of pre-generated scaffolds into the RIF to identify favorable ligand placements, we developed a physics-based generative approach within our ProBuilder^32,33^ protein design package that enables the de novo construction of protein backbones tailored to the target ligand. Our methodology is centered on helical protein architectures, which are prevalent in natural small molecule-binding proteins (e.g., GPCRs and nuclear receptors). Helical protein architectures are also frequently employed in de novo binder designs using docking approaches^17,18^, and particularly amenable to biosensor engineering. In each sampling trajectory, we initiate the process with an extended protein chain of the desired length, randomly assigning secondary structure elements in accordance with the target protein topology, such as four- or five-helix bundles.

To generate the folded protein structure, we begin by aligning a randomly selected residue to a randomly chosen RIF anchor point (**Fig. 1c**). This residue is designated as the folding root and remains fixed during the backbone generation process. Protein folding then proceeds outward from this root via Monte Carlo fragment assembly, utilizing a curated library of fragments derived from native structures. At each step, candidate moves are evaluated using a composite scoring function that simultaneously optimizes both protein folding and ligand binding. Protein folding is assessed by the residue-pair transform (RPX) score^34^, whereas ligand-binding is rapidly estimated based on the RIF. To enhance ligand specificity, privileged interactions, such as critical hydrogen bonds, can be upweighted during sampling. Monte Carlo moves are accepted or rejected according to standard Metropolis criteria, and sampling continues until convergence. To ensure thorough exploration of the conformational landscape, if a predefined threshold of consecutive unsuccessful moves is reached, the folding root is reassigned to another residue with a corresponding RIF anchor, thereby allowing all regions of the protein to sample alternative spatial relationships to the ligand (**Supplementary Fig. 1**). This generative, RIF-anchored protocol enables efficient sampling of global protein topologies optimized for ligand binding.

Backbones exhibiting well-folded structures and favorable ligand interactions (**Fig. 1d**) were subjected to sequence optimization using Rosetta^35^ and ProteinMPNN^36^. The resulting designs were rigorously filtered based on criteria including apoprotein folding, ligand interaction energy, the number and geometry of protein–ligand hydrogen bonds (or, in the case of metals, the quantity and geometry of coordinating residues), and overall shape complementarity. Additionally, the structural accuracy of the final models was assessed by comparing each design to AlphaFold2-predicted structures^37^. The selected designs encompass a diverse array of configurations (**Supplementary Fig. 2**), each featuring well-folded protein backbones and customized ligand-binding pockets precisely contoured to their respective targets.

### Design of small molecule binding proteins

To validate the efficacy of our ligand-binding design pipeline, we first applied our approach to three structurally distinct small molecules: dolutegravir (DTG, an antiviral drug targeting HIV; **Fig. 2a**), cianidanol (CIA, an antioxidant flavonoid; **Fig. 2b**), and DFHBI (a fluorogenic chemical dye; **Fig. 2c**). For each ligand, we generated helical bundle binders comprising 120 amino acids, consisting of five or six helices. For experimental characterization, we selected 96 DTG designs, 94 CIA designs, and 32 DFHBI designs, capturing a broad range of architectures (median pairwise backbone RMSD of 12.63 Å for DTG, 11.98 Å for CIA, and 12.68 Å for DFHBI, **Supplementary Fig. 2**). The majority of these proteins were successfully expressed in *E. coli* and were found to be soluble and monomeric by size exclusion chromatography. Ligand binding was assessed by isothermal titration calorimetry (ITC) for DTG and CIA binders. Initial screening of 96 DTG designs identified 14 binders (**Fig. 2a and Supplementary Fig. 3**), with the top candidate exhibiting an affinity of 4.94 μM. To further enhance binding affinity for DTG, we employed a helix-extension strategy by introducing two additional helices at either the N or C terminus to stabilize the binding helix and improve structural preorganization (**Supplementary Fig. 4a**). After evaluating 16 redesigned variants based on our top initial candidate, we achieved a 15.8-fold increase in affinity, reaching 971 nM (**Supplementary Fig. 4b,c**). For CIA, 20 out of 94 designs showed binding, with the highest measured affinity of 43.7 μM (**Fig. 3b and Supplementary Fig. 5**). For DFHBI, whose fluorescence is activated upon binding and rigidification by proteins, fluorescence assays revealed that 2 out of 16 designs produced a clear fluorescence increase following ligand incubation (**Supplementary Fig. 6**). The most effective binder demonstrated a measured affinity of 43.7 μM as determined by ITC (**Fig. 2c**). To confirm that ligand binding occurred within the designed pocket, we introduced mutations at key interface residues—either substituting contact residues with alanine or introducing bulky amino acids to disrupt binding through steric occlusion. ITC experiments on these mutants showed complete loss of ligand binding, supporting the intended binding mode (**Fig. 2**). Overall, these results demonstrate the high success rate and general applicability of our physics-based generative approach for the de novo design of small molecule-binding proteins.

**Fig. 2:**
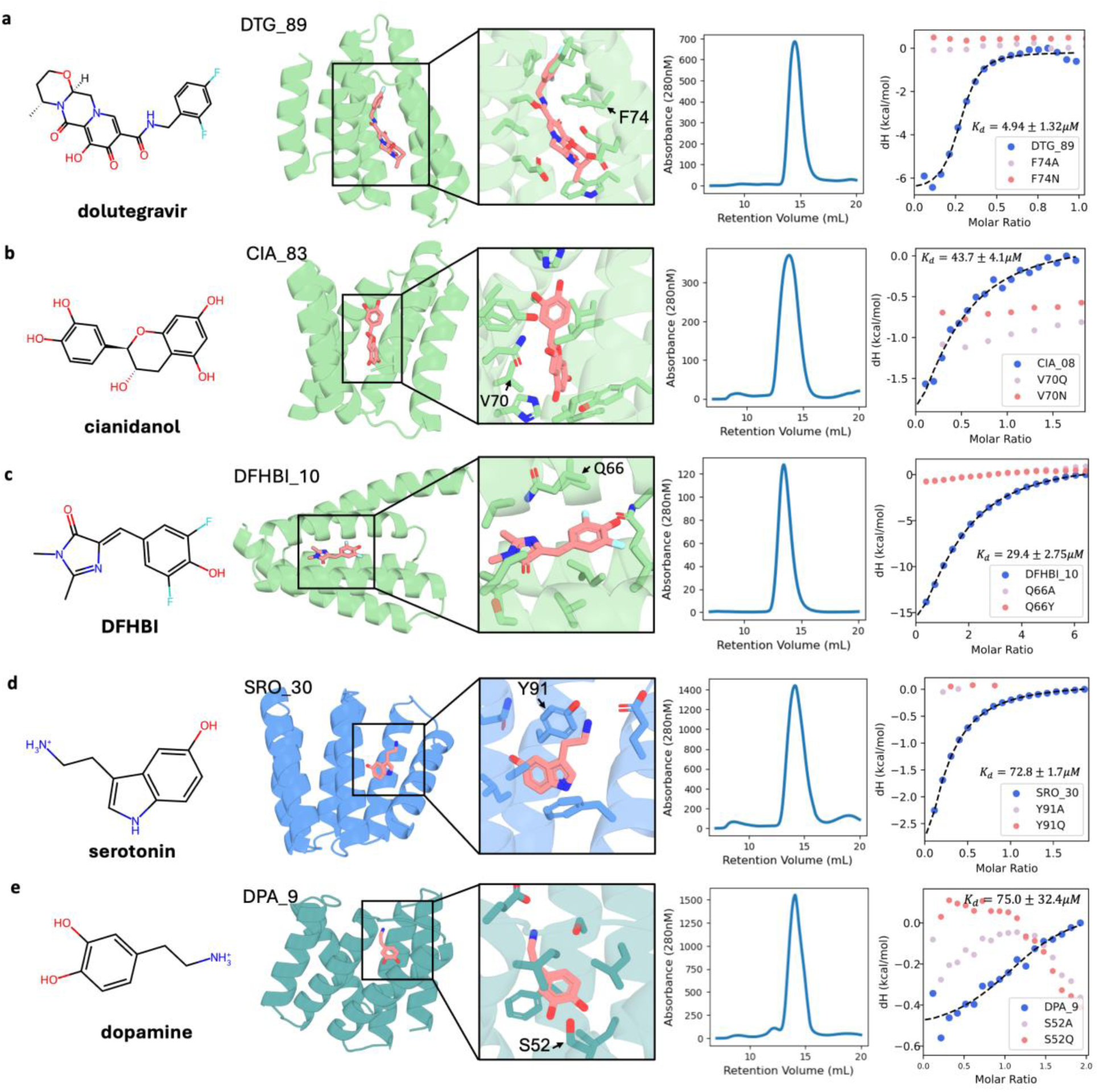
Experimental characterization of designed small-molecule binders. **a–e**, Experimental characterization of designed binders for DTG (**a**), CIA (**b**), DFHBI (**c**), SRO (**d**), and DPA (**e**). For each target, the chemical structure of the ligand is shown (left), followed by the design model of the protein–ligand complex. Zoomed-in views highlight pocket-binding residues, with the target residue for disruptive mutagenesis labeled. SEC traces confirm that the designed proteins exist as monomers, and ITC experiments validate binding. Mutations that disrupt interface residues abolished or significantly diminished binding in the pocket mutants, supporting the designed binding mode. See **Supplementary Figs. 4–7** for comprehensive data of all experimentally validated binders.

**Fig. 3:**
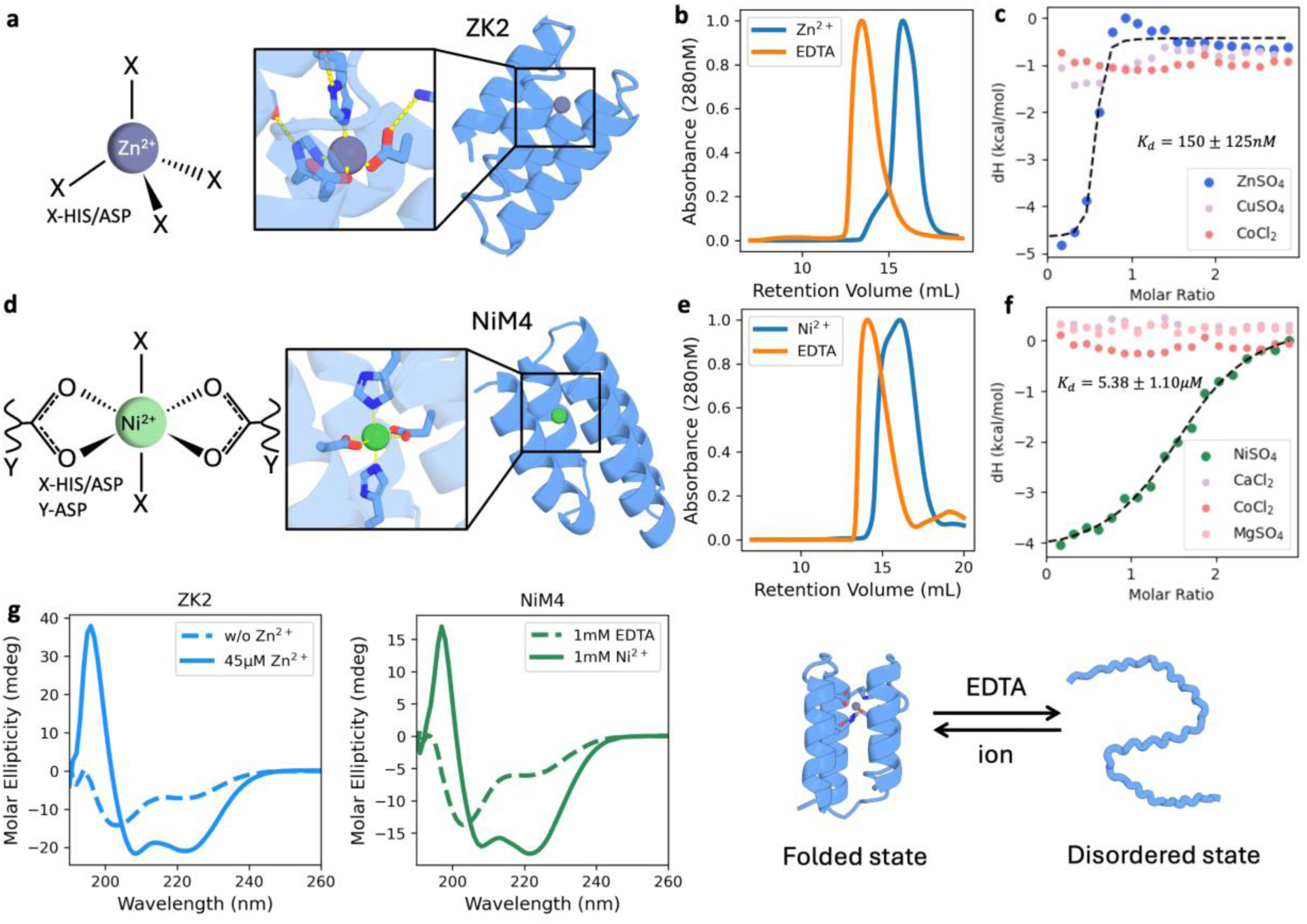
Experimental characterization of designed metal ion binders. **a,d**, Design models and zoomed-in views of the zinc binder ZK2 and the nickel binder NiM4. Zn²⁺ adopts a tetrahedral coordination geometry with four coordinating residues at each site. Ni²⁺ adopts an octahedral geometry, coordinated by two bidentate Asp residues and two monodentate His/Asp residues. **b,e**, SEC profiles show distinct elution differences between ion-bound (blue) and EDTA-treated (orange) samples, indicating an ion-dependent change in conformational compactness. **c,f**, Specificity and binding affinity were assessed by ITC. **g**, Circular dichroism spectra of ZK2 and NiM4 in the presence of target ions (solid lines) versus EDTA (dotted lines) demonstrate a transition from a disordered state (EDTA) to a folded state with defined secondary structure (ions). Right: Schematic illustration of the reversible conformational switch between the folded and disordered states driven by metal ion binding and chelation.

### Design of small molecule binding proteins targeting neurotransmitters

Previous efforts in small molecule binder design have predominantly targeted large, hydrophobic, or polar ligands with relatively high molecular weight^13,24,38,39^. Such large molecules offer more atomic contacts for favorable interactions, increasing both the likelihood and maximal affinity of generating effective binders. In contrast, engineering protein binders for very low molecular weight small molecules presents a substantial challenge, as these ligands provide fewer atoms for productive interactions^40,41–43^. For very small and polar ligands, successful binding demands exceptionally high shape and chemical complementarity to achieve sufficient binding energy. Our physics-based generative design approach enables precise tailoring of binding pockets, allowing us to target even very small and polar molecules. To rigorously test this capability, we selected serotonin (SRO) (**Fig. 2d**) and dopamine (DPA) (**Fig. 2e**). These two targets are low-molecular-weight neurotransmitters that are central to human biology and of considerable interest for biosensor development. Using our generative pipeline, we designed 120-residue, six-helix bundle scaffolds for each ligand and selected 38 SRO and 33 DPA designs for experimental characterization. The majority of the designs were successfully expressed in *E. coli* and ITC experiments revealed that six serotonin binders exhibited binding, with the highest measured affinity reaching 72.8 μM (**Fig. 2d and Supplementary Fig. 7**). For dopamine, which is even smaller and more polar ligand, we successfully identified one binder with an affinity of 75.0 μM (**Fig. 2e**). The low success for DPA highlighted the increased challenge of targeting such small and polar molecules. Importantly, interface-disrupting mutations abolished ligand binding in these proteins, confirming that the observed interactions were mediated by the designed binding pockets (**Fig. 2**).

To improve binding affinity, we performed in silico sequence optimization coupled with experimental screening on SRO_26 and SRO_30. In each optimization round, sequence variants were generated using Rosetta sequence design and LigandMPNN^44^ focused on ligand neighboring residues (**Supplementary Fig. 8a**). Following rigorous computational filtering, selected variants were experimentally characterized, with the highest-affinity binder advanced to the next optimization round. After four rounds of iterative optimization, we obtained a SRO_26 variant with a 15.7-fold improvement, achieving an affinity of 29.9 μM (**Supplementary Fig. 8b,c**). Notably, the accumulated mutations predominantly localized to the ligand-binding pocket and were primarily hydrophobic (**Supplementary Fig. 8b**), likely contributing to enhanced hydrophobic packing with the ligand. To our knowledge, SRO and DPA represent some of the smallest small-molecule ligands for which de novo protein binders with low micromolar affinity have been achieved. Collectively, these results underscore the generative power of our approach in creating highly complementary binding pockets, even for challenging small, polar molecules.

### Design of metal ion binding proteins

To demonstrate the generalizability of our approach for ligand binder design, we next applied our methodology to create proteins capable of binding zinc in a tetrahedral geometry (**Fig. 3a**) and nickel in an octahedral geometry^31^ (**Fig. 3b**). Histidine and aspartate/glutamate residues were utilized as coordinating residues, and two bidentate coordination residues were employed for nickel. For metal ions, we designed compact four-helix bundle proteins comprising 65 amino acids, with the coordinating residues and metal ions deeply buried at the protein core. The use of such small scaffolds was intended to ensure that, in the absence of the metal ion, the presence of unsatisfied polar coordination residues would destabilize the core and promote an unfolded state. Upon metal binding, the protein folds into the designed structure, thereby coupling ligand recognition to a global conformational change—an ideal property for metal ion sensor development. For each metal ion, we selected 10 designs for experimental characterization. His- tagged proteins were expressed in *E. coli*, purified using immobilized metal ion affinity chromatography (IMAC), and cleaved to remove the His-tag. SEC in the presence of the target metal showed that eight zinc and eight nickel binders eluted as monomers. SEC analyses conducted in the presence of EDTA, which chelates and removes metal ions, revealed clear shifts in the elution profiles for four zinc designs and one nickel design, indicating the formation of species with larger hydrodynamic radii (**Fig. 3a–b and Supplementary Fig. 9a**). These findings indicate that metal binding is indeed coupled to protein folding, as intended. Circular dichroism (CD) spectroscopy further supported this: in the presence of metal ions, the proteins displayed characteristic helical spectra, while removal of metal ions resulted in spectra indicative of unfolded structures (**Fig. 3c and Supplementary Fig. 9b–e**). ITC was used to assess binding affinity and selectivity. For zinc, the design ZK2 demonstrated a 1:1 protein-to-metal stoichiometry and a dissociation constant of 150 nM, with no detectable binding to Cu^2+^, Co^2+^, or Ni^2+^, indicating high specificity. This strong selectivity is particularly notable given the Irving–Williams series^42,45^, which typically favors Cu²⁺ over Zn²⁺ binding, yet our design successfully circumvents this thermodynamic preference to achieve exclusive zinc coordination.. Similarly, the nickel binder NiM4 displayed 1:1 stoichiometry with a Kd of 5.38 μM and showed no binding to Ca^2+^, Co^2+^, or Mg^2+^. These results demonstrate that our generative approach yields metal ion-binding proteins with high selectivity and specificity for their intended targets.

### Structure characterization

To validate both the overall architecture and ligand-binding modes of our designs, we determined high-resolution crystal structures of selected designs in complex with their target ligands. We successfully solved the structure of SRO_30 bound to serotonin at a resolution of 2.58 Å (**Fig. 4a and Supplementary Table. 7**). The backbone conformation closely matched the design model, with a Cα RMSD of 0.97 Å. Serotonin was well resolved in the electron density, and its configuration aligned closely with the design prediction (ligand RMSD of 1.997 Å using the protein backbone as reference). The hydrophobic indole ring of serotonin was tightly packed against neighboring hydrophobic residues, while its amine group formed polar interactions with serine and glutamate side chains. We further determined the crystal structure of the L7F mutant of SRO_26 (**Fig. 4b**). The structure was resolved at 2.00 Å, and the overall conformation matched the design model with atomic accuracy. Notably, in SRO_26_L7F, the bound serotonin adopted a flipped orientation relative to the original design, which might be attributable to the sequence optimization process. Additionally, we solved the structure of DPA_9 in complex with dopamine at 2.47 Å resolution (**Fig. 4c**). The protein backbone exhibited atomic precision relative to the design (Cα RMSD of 0.791 Å), and the dopamine ligand was clearly resolved. The configuration of the ligand was accurately recapitulated, engaging in designed hydrophobic and polar contacts, with a ligand RMSD of just 0.883 Å. The crystal structure of ZK2 complexed with zinc was determined at 1.23 Å resolution (**Fig. 4d**). The protein backbone showed exceptional agreement with the design (Cα RMSD of 0.446 Å), and both the zinc ion and its four coordinating residues were observed as predicted. Overall, the high level of concordance between the crystal structures and our computational models underscores the accuracy and robustness of our physics-based generative ligand-binding protein design approach.

**Fig. 4:**
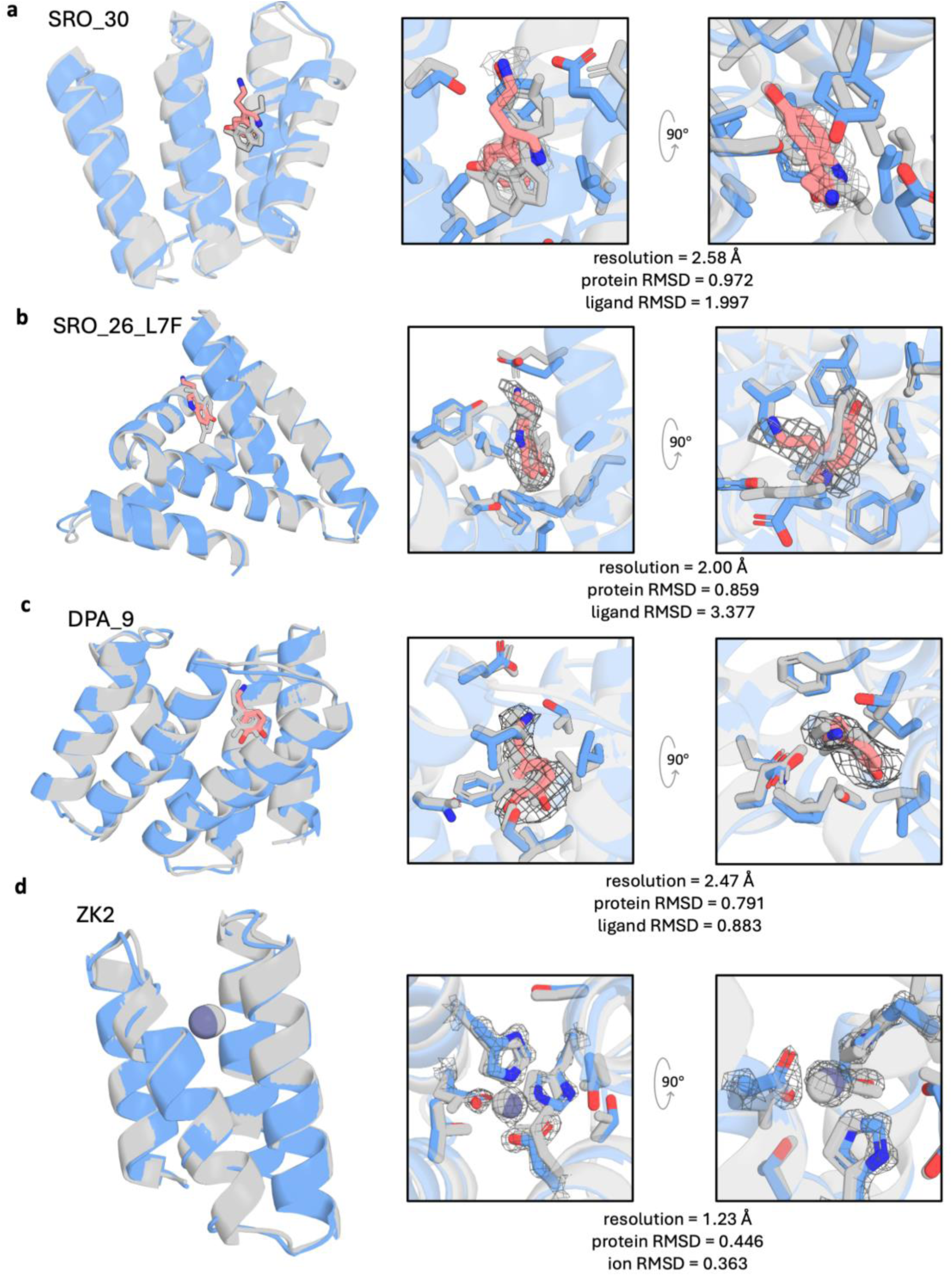
X-ray crystallographic structures confirm the high design accuracy. Comparison of computational design models (gray) with experimental crystal structures (blue) for SRO_30 (**a**), SRO_26_L7F (**b**), DPA_9 (**c**), and ZK2 (**d**). Composite omit electron density of the ligand and neighboring residues are contoured at 0.5σ(SRO_26, SRO_30, DPA_9), 2.0σ(ZK2).

### Developing neurotransmitter sensors

Our de novo designed small molecule-binding proteins feature simple helical bundle topologies, which facilitate engineering and make them well suited for biosensor development. To demonstrate their potential, we focused on biosensors for SRO and DPA, both central to neural signaling and direct monitoring of neural activity^46,47,48^. For these targets, we employed a split protein strategy: each ligand-binding protein was divided into two fragments and fused to the termini of a circularly permuted superfolder GFP (cpsfGFP^49^) (**Fig. 5a**). In the presence of their respective neurotransmitter, ligand binding promotes the re-association of the protein fragments, which can alter the environment of the cpsfGFP chromophore, leading to a fluorescence signal change^5^. SRO_30 is a six-helix bundle protein. We split this protein into two parts at various loop regions, generating five different fragment combinations. For each pair of fragments, we created error-prone libraries using error-prone PCR and fused the fragments to the N- and C-termini of cpsfGFP. To identify functional SRO sensors, we utilized a plate-based high-throughput screening strategy to select variants with beneficial mutations and optimal splitting sites (**Supplementary Fig. 10**). Colonies were screened using a plate-based fluorescence assay, with fluorescence measured before and after serotonin exposure. Colonies displaying robust fluorescence responses following serotonin exposure were selected for further analysis. Through this screening, we identified a variant termed wcSRO_30_0.1, which exhibited a ΔF/F₀ of 27.6% in response to 10 mM serotonin (**Supplementary Fig. 10c**). Colony sequencing revealed that this variant was generated by splitting five helices at the N terminus and one helix at the C terminus (**Fig. 5A**). Notably, it contains an eight-amino-acid helical repeat, copied from the C-terminus of the split N-terminal fragment, which is inserted immediately after the cleavage site within the C-terminal fragment. This duplicated helix region is most likely introduced during PCR or gene assembly. We hypothesized that this repeat might increase the energy barrier for self-association between the split parts, reducing their propensity for spontaneous reconstitution and thereby conferring SRO sensitivity (**Supplementary Fig. 11**). Building on this observation, we further screened interface variants by replacing large hydrophobic residues at the split interface with smaller ones to further diminish self-association propensity. This led to the identification of a variant, wcSRO_30_0.5, which displayed a much greater ΔF/F₀ fold change of 179.4% in response to 10 mM serotonin. Finally, we explored various linker combinations connecting the split fragments to cpGFP, ultimately identifying a final variant, wcSRO30_1.0, which showed a ΔF/F₀ of 339.9% (**Fig. 5b and Supplementary Fig. 10**).

**Fig. 5:**
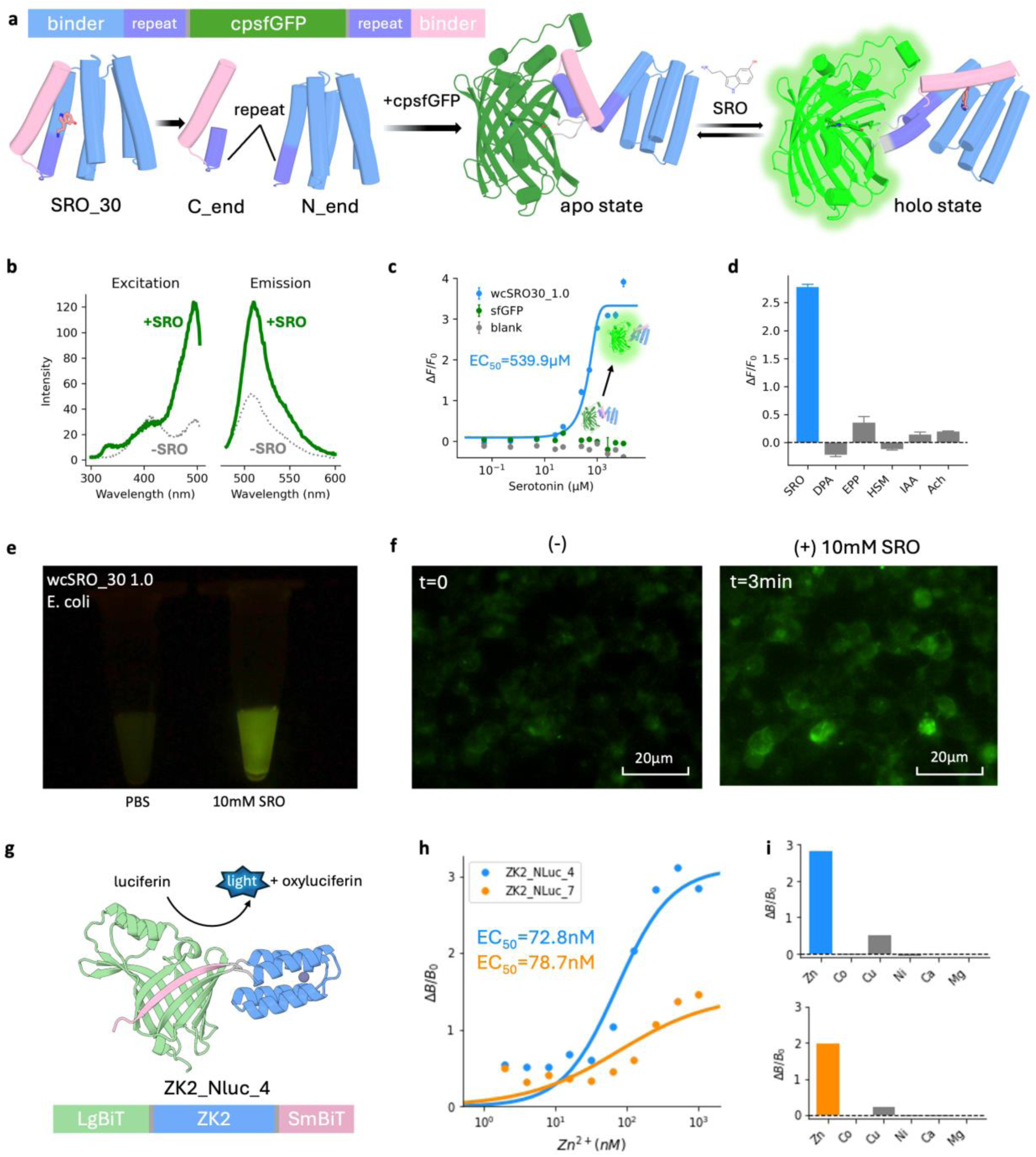
Development of functional biosensors from de novo designed ligand binders. **a**, Schematic illustrating the development of the serotonin sensor wcSRO30_1.0. SRO_30 was split into two fragments and fused to the termini of cpsfGFP. High-throughput screening identified a repeat helix variant, which likely reduces self-association. Further optimization—through interface destabilization and linker engineering—resulted in wcSRO30_1.0. **b**, Excitation and emission spectra of wcSRO30_1.0 in the presence (green) and absence (gray) of 10 mM serotonin. **c**, Fluorescence titration of wcSRO30_1.0, demonstrating a dose-dependent response to serotonin. **d**, Specificity profile of wcSRO30_1.0 against structurally similar ligands (1 mM each): dopamine (DPA), epinephrine (EPP), histamine (HSM), indole-3-acetic acid (IAA), and acetylcholine (ACh). **e**, *E. coli* cells expressing wcSRO30_1.0 exhibit a clear fluorescence signal change upon incubation with 10 mM SRO, visualized under 470 nm blue light. **f**, Fluorescence microscopy images of HEK293T cells expressing wcSRO30_1.0 before (t = 0) and after (t = 3 min) the addition of 10 mM serotonin, demonstrating SRO response in mammalian cells. **g**, Schematic of the de novo zinc sensor development. Based on the zinc binder ZK2, the sensor uses a split NanoLuc luciferase complementation strategy, fusing binder termini to LgBiT (light green) and SmBiT (light pink). **h**, Bioluminescence dose-response curves for the ZK2_NLuc4 and ZK2_NLuc7 variants, showing nanomolar sensitivity to Zn²⁺. **i**, Ion selectivity profile of the zinc sensors tested against various metal ions (1 μM), demonstrating high specificity for Zn²⁺ over Co²⁺, Cu²⁺, Ni²⁺, Ca²⁺, or Mg²⁺.

Our success with the SRO sensor suggests that, since the designed binders are composed of well-packed regular helix bundles with hydrophobic residues, reducing the self-association propensity of the split fragments is critical for developing functional sensors. This can be achieved by introducing repeat helices to block self-association, implementing interface mutations to reduce binding energy. To test the generalizability of these principles, we designed a new SRO sensor based on a different binder, SRO_26. We split the protein into a 1-helix and a 5-helix configuration, copied residues from the adjacent helix onto the 1-helix fragment as a repeat helix and tested several repeat lengths. After fusing these split fragment variants with cpGFP, we quickly identified a variant with a ΔF/F₀ of 13.8%, corresponding to a repeat helix of 7 amino acids (**Supplementary Fig. 12**). To further decrease self-association, we truncated the 1-helix part and identified a variant with a five-residue deletion, achieving a ΔF/F₀ of 78.9%. The same principle was applied to the DPA sensor DPA_9, directly yielding a 1–5 split variant with a repeat of 7 amino acids and a ΔF/F₀ of 24.3% (**Supplementary Fig. 13**). Screening linker variants, we obtained a construct with a ΔF/F₀ of 37.5%, marking the first reported soluble dopamine biosensor. By strategically balancing the repeat regions and helix truncation, we effectively suppressed the high background noise caused by fragment self-association and increased ligand sensitivity. The SRO_26 and DPA_9 based sensors are all identified through low throughput screening (n = 24 and n = 9 for SRO_26 and DPA_9 respectively) without hight throughput screening and directed evolution. Although their current performance is suboptimal, they provide a promising foundation for further optimization toward ultra-high performance in real-world applications.

### Characterization of the biosensor

We further characterized of the property and performance of the SRO biosensor wcSRO30_1.0 under a variety of conditions. In the absence of serotonin, wcSRO30_1.0 exhibited excitation/emission maxima at 417/507 nm (**Fig. 5b**). Upon serotonin addition, these maxima slightly shifted to 493/512 nm, indicating a alteration in the chromophore environment upon ligand binding. Compared with the well-established iSeroSnFR sensor^47^, wcSRO30_1.0 exhibited improved ligand sensitivity (EC₅₀ = 539.9 μM vs. 959 μM for iSeroSnFR), while the dynamic fluorescence response remains lower ( ΔF/F₀ = 411% vs. 930% in *E. coli*). ITC measurements showed a binding affinity of 105 μM (**Supplementary Fig. 10e**). Specificity assays confirmed that wcSRO30_1.0 selectively responds to serotonin, with minimal cross-reactivity observed for five structurally similar ligands (dopamine, acetylcholine, epinephrine, auxin, and histamine) at 1 mM concentrations (**Fig. 5d**). Notably, the SRO sensing module in wcSRO30_1.0 is significantly smaller than that of iSeroSnFR (129 a.a. vs 272 a.a), which may offer advantages in applications with limited delivery system capacity. *E. coli* cells expressing wcSRO30_1.0 exhibited a SRO concentration dependent fluorescence change after incubation with SRO and demonstrated high reversibility following SRO washout (**Fig. 5e, Supplementary Fig. 10g and Supplementary Fig. 14a**). When expressed in HEK293T cells, wcSRO30_1.0 exhibited a clear, concentration-dependent fluorescence change in response to SRO as measured by flow cytometry (**Supplementary Fig. 14b**). Real-time fluorescence imaging further validated these findings, showing a pronounced increase in fluorescence following a three-minute exposure to 10 mM serotonin (**Fig. 5f**).

### Developing metal ion sensors

Our metal ion-binding proteins are distinguished by a deeply buried metal coordination site within the protein core, with metal ion binding tightly coupled to protein folding (**Fig. 3c**). We leveraged this property for biosensor development using a split-enzyme reconstitution strategy. Specifically, the two fragments of NanoLuc luciferase^50,51^ were fused to the N- and C-termini of the zinc-binding protein ZK2 (**Fig. 5g**). In the absence of zinc, the protein remains largely unfolded and extended, which keeps the split NanoLuc fragments apart and results in minimal luminescence. Upon zinc binding, the protein adopts its designed fold, bringing the N- and C-termini—and thus the NanoLuc fragments—into close proximity, enabling reconstitution of enzyme activity and generation of a luminescent signal.

We tested a series of linker variants and identified two constructs, ZK2_Nluc_4 and ZK2_Nluc_7, that exhibited robust increases in luminescence upon zinc addition. Titration experiments revealed EC₅₀ values of 72.8 nM and 78.7 nM (**Fig. 5h**), consistent with binding affinities measured by ITC. Specificity assays confirmed high zinc selectivity for the sensors, which generate high luminescence in the presence of zinc ions. Copper induced minimal signal, presumably due to its potent electrophilicity, whereas all other metal ions produced no measurable response (**Fig. 5i**). The precise architecture of the metal coordination sphere is critical for conferring this selectivity. These results demonstrate that our design strategy, which tightly couples metal ion binding with protein folding, enables highly specific metal ion sensing and highlights the power and versatility of our generative approach for engineering sensitive and selective metal ion biosensors.

## Discussion

Our physics-based generative approach provides a robust and versatile framework for the de novo design of small molecule and metal ion binders. By constructing protein architectures directly around the shape and chemical properties of the target ligand, our method eliminates the reliance on pre-existing scaffold libraries required for physics-based docking and enables precise customization of binding pockets. This bottom-up strategy allows the design of binding proteins tailored to a broad range of targets, including highly challenging small and polar small molecule ligands. Compared to recent deep learning-based generative methods^22,23,24,25,26^, our approach affords explicit control over protein topology throughout the scaffold generation process, facilitating the construction of structures that are optimally suited for biosensor engineering. The employment of simple, well-defined helical bundle scaffolds further enhances modularity and engineerability, as these architectures are ideally suited for strategies such as protein splitting, enabling direct coupling of ligand binding to protein association. In addition, our platform explicitly models both protein folding energetics and protein–ligand interaction energies, providing direct control over the stability and binding landscape of the designed proteins. This allows for the creation of proteins that are intrinsically unstable in the absence of ligand, yet adopt a well-folded, low-energy conformation upon ligand binding. In contrast to previous methodologies focused primarily on static, high-affinity binding, our strategy enables direct coupling of ligand recognition to conformational transitions, providing a robust mechanism for signal transduction in biosensor applications.

Despite significant advances in de novo protein design, engineering functional biosensors remains a complex and challenging task. We demonstrate the practical utility of our binder designs for biosensor development through two distinct strategies. For SRO and DPA binders, the modularity of helical bundles enabled successful splitting into separate fragments, which were fused to cpGFP to generate highly specific sensors. A major challenge with split protein sensors is potential self-association of fragments in the absence of ligand, resulting in elevated background signals and reduced sensitivity. We identified a fragment duplication strategy—duplicating a specific protein segment to increase the energetic barrier for self-association—which enables direct engineering of ligand-induced sensor activation. Additionally, we demonstrated that interface-weakening mutations and truncations can further enhance ligand responsiveness. Furthermore, the ligand-induced folding approach provides a straightforward platform for biosensor construction. By coupling ligand binding to protein folding, the resulting conformational change can be readily transduced to enzyme reporters or FRET pairs, enabling robust signal output. Our generative design methodology not only broadens the repertoire of ligand binders, but also offers efficient routes for engineering responsive biosensors. Although the current sensors exhibit promising performance, there remains considerable potential for further optimization. Targeted mutagenesis and directed evolution could further enhance ligand sensitivity and response dynamics, expanding their utility across a broader range of biological and biomedical applications.

In summary, our physics-based generative approach provides a versatile platform for the de novo design of ligand binders and biosensors, offering precise control over the structural and energetic features that underlie ligand-induced signal transduction. As this methodology is further refined, it has the potential to facilitate the development of biosensors with enhanced sensitivity and selectivity. Looking forward, integration with advances in deep learning, high-throughput screening, and in vivo validation may further expand the range and diversity of accessible sensing modalities. We anticipate that the tools and design strategies presented here will contribute to the real-time monitoring of complex biological processes and improved disease diagnostics.

